# Chemical composition, antimicrobial and antioxidant activities of essential oils from the receptacle of sunflower (*Helianthus annuus* L.)

**DOI:** 10.1101/2020.07.21.213587

**Authors:** Xinsheng Liu, Xinlu Li, Zian Qiao, Wannan Li, Bo Gao, Lu Han

## Abstract

Sunflower (*Helianthus annuus* L.) contains active ingredients, such as flavonoids, alkaloids and tannins. However, few studies have focused on essential oils from the receptacle of sunflower (SEOs). The chemical composition, antimicrobial and antioxidant activities of SEOs were assessed whether SEOs could be used as nature preservatives and medicines. The extraction rate of SEOs was 0.4% by using hydro-distillation and solvent with petroleum ether, which is much higher than that by only using hydro-distillation (0.2%) in the previous study. A total of 81 volatile constituents of SEOs were identified by gas chromatography-mass spectrometry (GC-MS). The main constituents of SEOs were α-pinene (31.17%), calamenene (5.66%), α-terpineol (4.71%), verbenone (3.15%), kaur-16-ene (2.52%) and terpinolene (1.47%). The antimicrobial activities of SEOs were assessed against three bacteria (*E. coli*, *P. aeruginosa*, and *S. aureus*) and two fungi (*S. cerevisiae* and *Candida albicans*). The MIC of SEOs against *P. aeruginosa* and *S. aureus* was 0.2 mg/mL. The MIC of SEOs against *S. cerevisiae* was 3.2 mg/mL. The MIC of SEOs against *E. coli* and *Candida albicans* was 6.4 mg/mL. The results showed that SEOs had high antibacterial and antifungal actions. The antioxidant activity was determined with three different analytical assays (DPPH, ABTS and iron reducing ability). The results of antioxidant activities showed that SEOs had high antioxidant activities. The results proved that SEOs could be used as natural preservatives and medicines, due to its excellent antimicrobial and antioxidant activities.

## Introduction

Sunflower (*Helianthus annuus* L.) is an annual dicotyledonous plant in the family *Asteraceae*, which is widely distributed across North America, Eastern Europe and Northern China for production of oils, seeds and snacks [1]. Sunflower roots, stems, leaves and seeds contain many active ingredients, including phenols, flavonoids, alkaloids, active proteins, and tannins, which have antibacterial, anti-inflammatory, antioxidant, antipyretic, and diuretic effects [2].

Essential oils as important natural active substances exist in roots, leaves and fruits of plants, such as *Citrus reticulate* and *Origanum vulgare* [3, 4], which have been widely used as natural antibacterial and antioxidant ingredients in pharmaceutical and food industries [5, 6, 7]. The main chemical component of essential oils from industrial hemp (*Cannabis sativa* L.) and mediterranean herbs was terpenoid, which had great antioxidant and antibacterial activities [8, 9]. The chemical component of essential oils from flowers and leaves of sunflowers has been analyzed in previous research [10, 11]. However, the essential oils from the receptacle of sunflower have not been reported. It proved that the receptacle of sunflower reduced the level of uric acid, which may be used as a Chinese traditional drug for treating anti-gout [12]. Few studies have forced on the composition and biological activities of essential oils from the receptacle of sunflower.

The aim of this study was to evaluate the chemical composition, antimicrobial and antioxidant activities of SEOs, in order to determine the potential use of the receptacle from sunflower in food and medical filed. The SEOs were extracted by hydro-distillation and solvent method with petroleum ether. The chemical composition of SEOs was identified by GC-MS. Antimicrobial activities of SEOs was assessed against three bacteria (*E. coli*, *P. aeruginosa*, and *S. aureus*) and two fungi (*S. cerevisiae* and *Candida albicans*) by using a microplate reader with 96-well microplate. The antioxidant activity was determined with three different analytical assays (DPPH, ABTS and iron reducing ability). The chemical composition and biological activities of SEOs would provide scientific guidance for developing and utilizing SEOs.

## Materials and methods

### Chemical reagents and solvents

Tetracycline hydrochloride, miconazole nitrate, 2,2-diphenyl-1-picrylhydrazyl (DPPH) and (*±*)-6-hydroxy-2, 5, 7, 8-tetramethylchromane-2-carboxylic acid (Trolox) were purchased from Source Leaf Biology Co. (Shanghai, China). Potassium persulfate and 2, 2′-azino-bis (3-ethylbenzothiazoline-6-sulfonic acid) (ABTS) were purchased from Meilun Biology Co. (Dalian, China). Iron ion reduction capacity kit was purchased from Congyi Biology Co. (Shanghai, China). All other chemical reagents and solvents were of analytical grade.

### Extraction of essential oils from the receptacle of sunflower

The receptacles of sunflower (*Helianthus annuus* L.) were collected in Baicheng (Jilin, China). The essential oils were extracted by using both hydro-distillation and organic solvent method, according to modified protocol of Dong and Lawson [13, 14]. The powder of the receptacle from sunflower was soaked in saturated sodium chloride solution for 2 h. SEOs were extracted with petroleum ether by hydro-distillation method for 10 h in the volatile oil extractor. The extracts were evaporated to dryness using a Rota-vapor. SEOs were stored at −20 °C for use.

### Analysis of chemical compositions of essential oils

Analysis of chemical compositions of SEOs was performed using Agilent 5975 (Agilent Technologies, Santa Clara, CA, USA). Helium was used as a carrier gas at 20 mL/min. The oven column temperature was programmed 60 °C, initial hold time of 3 min, to 240 °C at 10 °C/min with final hold time of 15 min, using helium as carrier gas at a flow rate of 20 mL/min. The injector and ion source temperatures were 260 °C. Use Agilent 19091N-133 capillary column (Polyethylene Glycol, 30 mm × 0.25 mm internal diameter; film thickness 0.25 μm). Comparing the retention indices, the retention index was calculated for the same series of normal paraffins using the arithmetic index, and the mass spectrum was compared with the mass spectrum reported in NIST17 to identify the chemical compositions [15].

### Antimicrobial effects

Microbial cultures of three bacterial strains were *E. coli* (ATCC 25922), *P. aeruginosa* (ATCC 15442), and *S. aureus* (ATCC 25923). Fungal strains were *Candida albicans* (ATCC 10231) and *S. cerevisiae* (ATCC 9080). All the microbial strains in this study were purchased from Huan Kai Microbiology Technology Co. (Guangzhou, China).

Bacteria stains were resuscitated in lysogeny broth (LB) solid medium as resuscitation fluid, which transferred to LB liquid medium and incubated at 37 °C for 24 h. The fungi were resuscitated in yeast extract peptone dextrose medium and adenine (YPDA) solid medium. Then, it was transferred to YPDA liquid medium and incubated at 26 °C for 48 h.

### Determination of antimicrobial activities

The minimum inhibitory concentration (MIC) and minimum bactericidal concentration (MBC) were measured by using the microplate reader (ELx800, BioTek, USA) [16]. Each monomer of essential oils (α-pinene, α-terpineol, terpinolene, and verbenone), monomer mixture with α-pinene, α-terpineol, terpinolene, and verbenone, and SEOs in the culture medium had nine different concentrations, which were 0.05 mg/mL, 0.1 mg/mL, 0.2 mg/mL, 0.4 mg/mL, 0.8 mg/mL, 1.6 mg/mL, 3.2 mg/mL, 6.4 mg/mL, and 12.8 mg/mL. The 96-well microplate were mixed with 179 μL culture medium, 20 μL sample, and 1 μL 2.0 × 10^6^ CFU/mL bacteria or 2.0 × 10^5^ CFU/mL fungi. The bacteria were cultured at 35-37 °C for 24 h, and the fungi were cultured at 25-26 °C for 48 h. The absorbance at 600 nm was measured by using the microplate reader. Tetracycline hydrochloride and miconazole nitrate were used as positive controls of antibacterial and antifungal experiments separately. Dimethyl sulfoxide (DMSO) (5%, W/V) and DMSO (1%, W/V) were used as negative controls of antibacterial and antifungal experiments separately. The experiments were carried out in triplicate.

### Determination of antioxidant activities

Three different analytical assays, including ABTS, DPPH, and FRAP were assessed the antioxidant activities of SEOs, because one antioxidant method cannot evaluate the antioxidant activity of SEOs accurately.

#### ABTS free radical scavenging assay

ABTS radical scavenging activity was according to the modified protocol of Kang [17]. The ABTS working solution was mixed with 2.6 mmoL K_2_S_2_O_8_ and 7.4 mmoL ABTS, which was incubated for 12 h at room temperature in dark and diluted 40-45 times with ethanol. We added 0.5 mL sample to 2 mL ABTS working solution, which was incubated for 6 min at room temperature in dark. The absorbance was measured at 734 nm. Trolox was used as a positive control.

The ABTS scavenging rate was determined by the following formula:

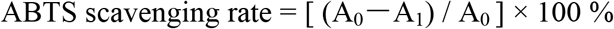

A_0_ was the absorbance of the negative control without SEOs and A_1_ was the absorbance of the test sample with SEOs.

#### DPPH radical scavenging

DPPH radical scavenging activity was determined according to modified protocol of Das method [18]. Ethanol and DPPH were mixed for preparing 0.08 mmoL/L DPPH solution, which was stored in dark. The 1 mL sample and 3 mL DPPH solution was mixed and conducted at room temperature for 30 min in dark. The absorption value was measured at 517 nm. Anhydrous ethanol and Trolox was as the negative control and positive control respectively.

The DPPH radical scavenging capacity was determined by the following formula:

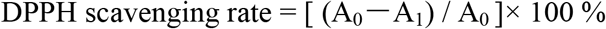

A_0_ was the absorbance of the negative control without the SEOs and A_1_ was the absorbance of the test sample with SEOs.

#### Determination of iron ion reducing assay

The iron ion reducing ability was determined by using Congyi Biology kit (Shanghai, China), according to the method of Prussian blue [19]. The reduction ability was determined by amount of Prussian blue Fe_4_ (Fe (CN) _6_). The antioxidant activity can reduce the ferric iron of potassium ferricyanide to ferrous ions to form Prussian blue. The material had a maximum absorption peak at 700 nm. The larger absorption value means the better antioxidant capacity of the sample.

### Statistical analysis

All the experiments were performed at least two independent times. Each time was conducted with three replications. One-way analysis of variance (ANOVA) and the mean comparisons were performed on all antimicrobial and antioxidant data using SPSS 20.0. Program (IBM Corporation, NY, USA). Duncan’s multiple range tests were used to calculate the mean values. Differences between mean values at P < 0.05 were considered significantly.

## Results and Discussion

### Extraction of SEOs

The extraction rate of SEOs was 0.4% by using both hydro-distillation and petroleum ether method, which was much higher than that (0.2%) by using only hydro-distillation method in the previous research [11]. Saturated sodium chloride was used to reduce the solubility of the substance in water for raising the extraction rate of essential oils [20]. Since the essential oil was continuously dissolved in the organic solvent during the extraction process, the reaction between the essential oil and water is reduced [13].

### Chemical compositions of SEOs

As shown in Table 1, A total of 81 compounds were identified by GC-MS. The major chemical composition was α-pinene with 31.17% of the total peak area. The other main components were calamenene (5.66%), α-terpineol (4.71%), verbenone (3.15%), kaur-16-ene (2.52%) and terpinolene (1.47%). Among of the 81 compounds, most of them were belonged to monoterpenoids (64.31%), mainly including α-pinene (31.17%), calamenene (5.66%), α-terpineol (4.71%), verbenone (3.15%) and terpinolene (1.47%). The other components of SEOs were belonged to sesquiterpenoids (15.5%), aromatics (10.26%), and diterpenoids (2.52%). The results of chemical compositions of SEOs showed that monoterpenoids (64.31%) were the predominant components of SEOs.

**Table 1.**
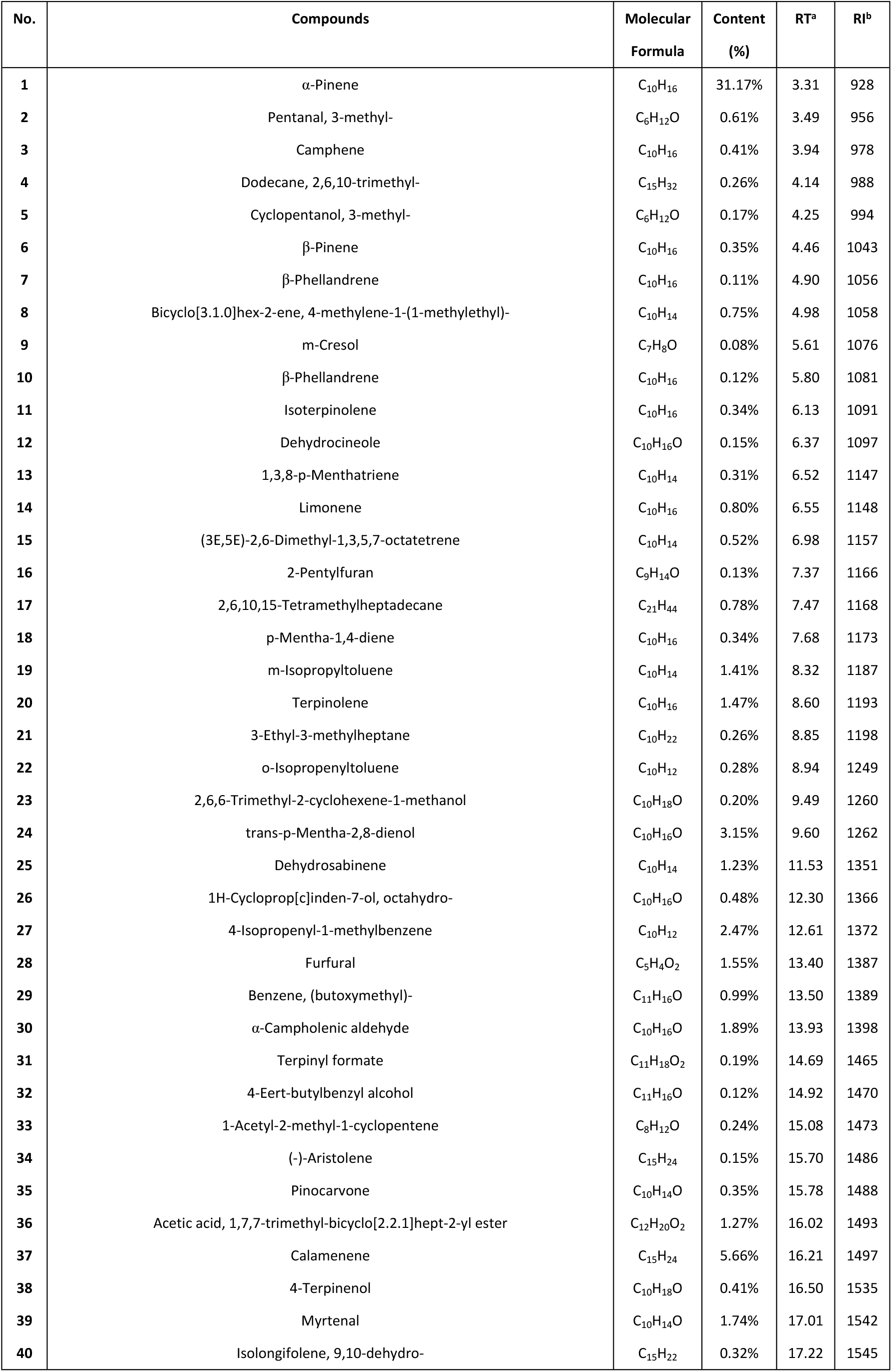
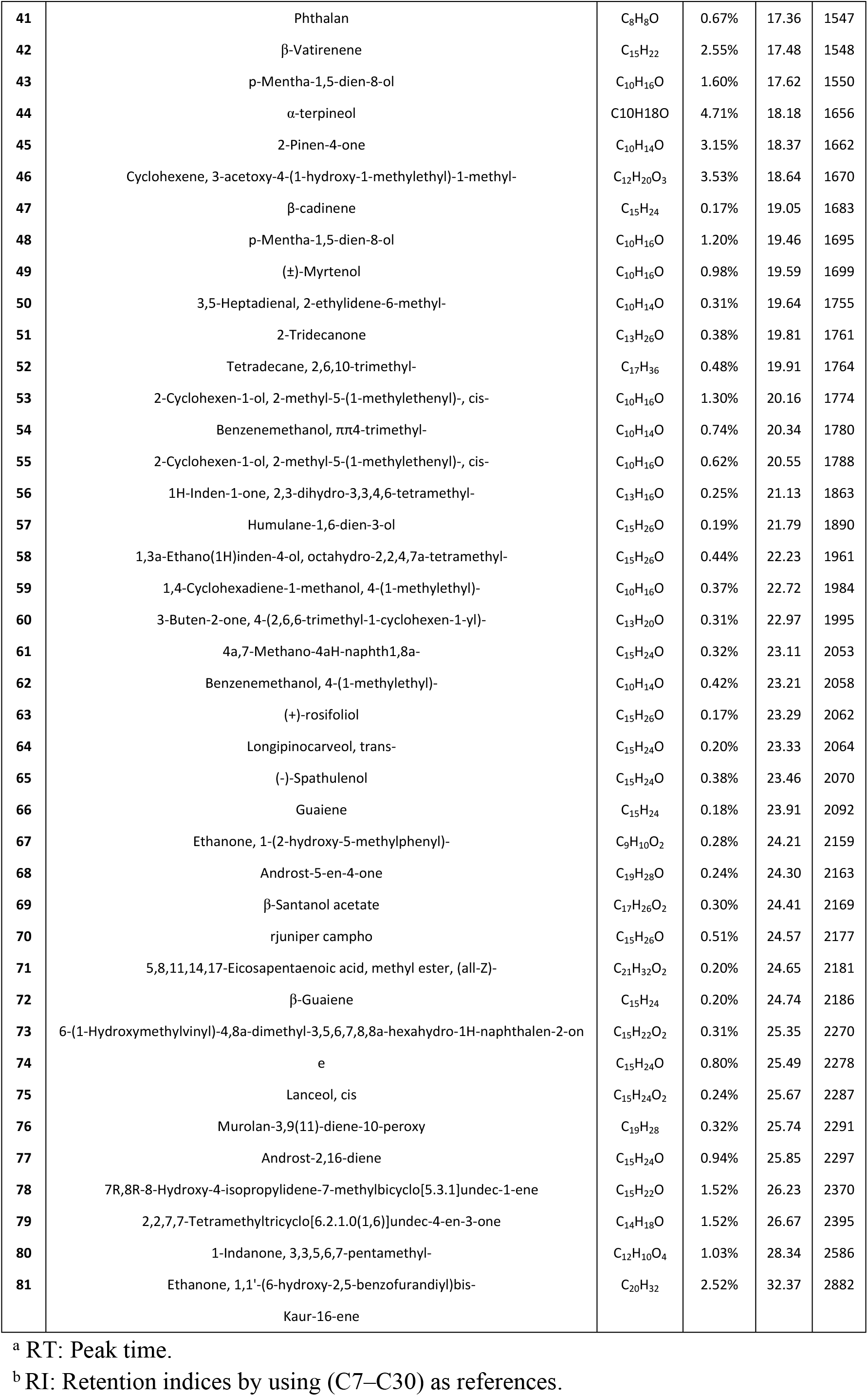
Chemical compositions of essential oils from the sunflower receptacle.

### Antimicrobial activities of SEOs against bacteria and fungi

Antimicrobial activities of SEOs were evaluated against bacteria (*E. coli*, *P. aeruginosa*, *S. aureus*) and fungi (*S. cerevisiae* and *Candida albicans*) using 96-well microplate. Tetracycline hydrochloride and miconazole nitrate were used as positive controls against bacteria and fungi separately.

#### Antibacterial effects

The antibacterial activities of SEOs and monomers were evaluated against *E. coli*, *P. aeruginosa* and *S. aureus* (Table 2 and S1Table). The results showed that the MIC of SEOs against *P. aeruginosa* and *S. aureus* was 0.2 mg/mL. The MIC of SEOs monomer mixtures against *P. aeruginosa* and *S. aureus* were 1.6 mg/mL. The MIC of α-pinene against *P. aeruginosa* and *S. aureus* were 6.4 mg/mL. The MBC of α-terpineol against *P. aeruginosa* and *S. aureus* were 6.4 mg/mL. The MIC and MBC of α-terpineol against *E. coli* were 6.4 mg/mL. Privious study showed that α-terpineol had antibacterial activity by changing the morphology of *E. coli* [21].

**Table 2.**
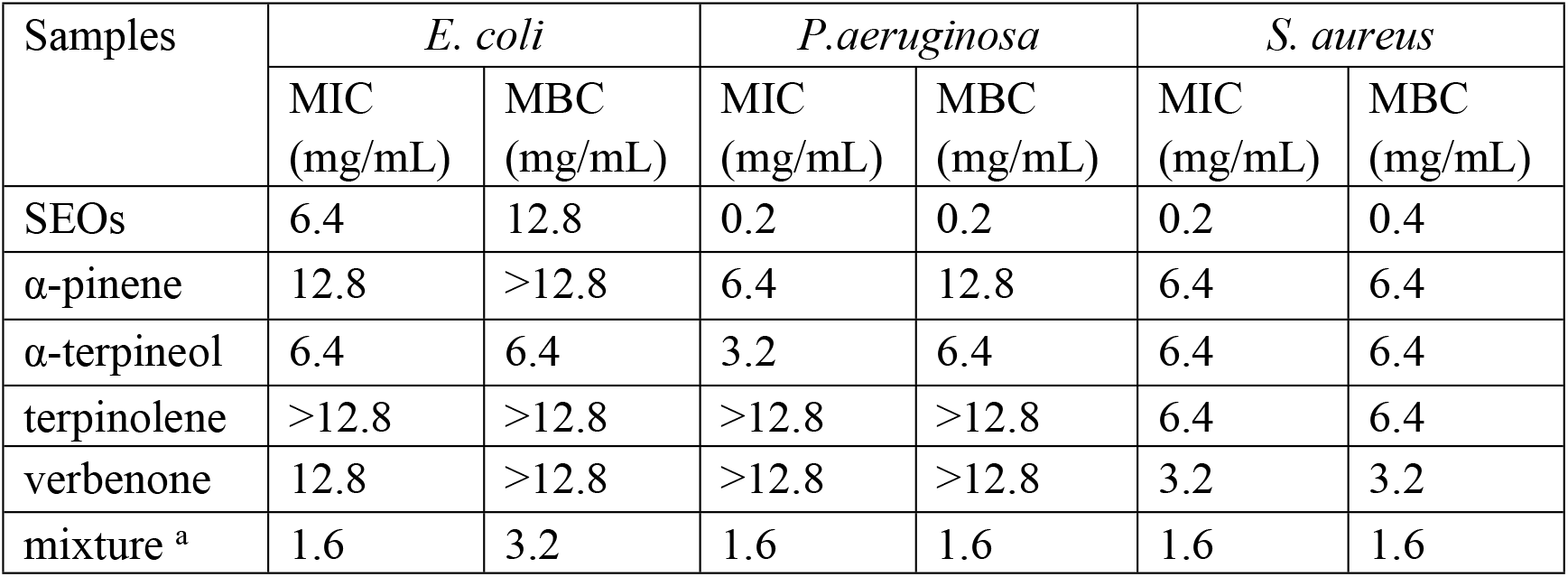
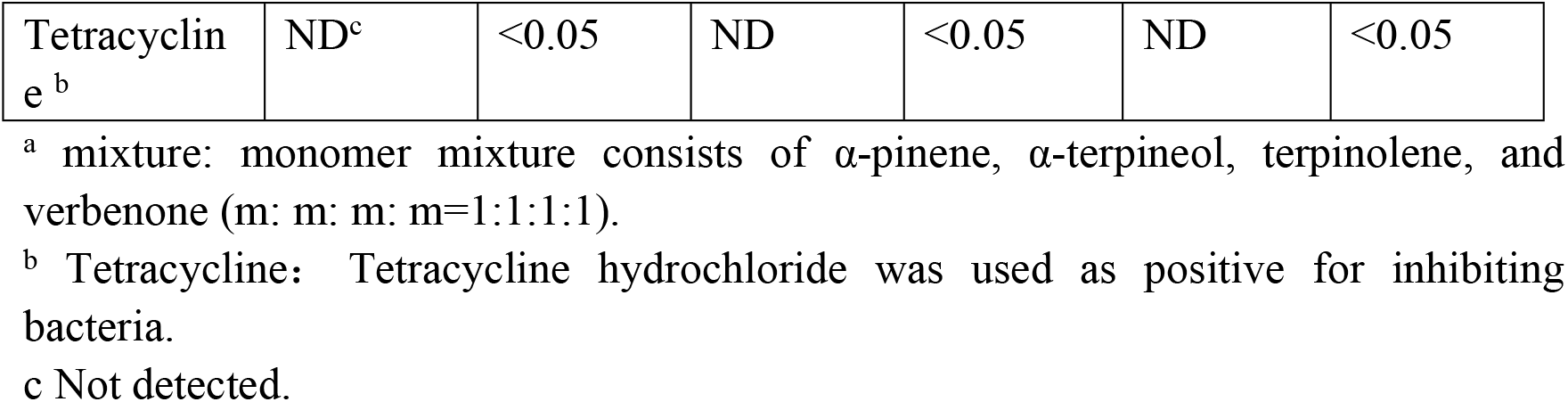
Antibacterial activities of SEOs and the dominant constituents.

The MIC and MBC of SEOs were lower than that of the monomers mixture and each monomer, which implied that the antibacterial effect of SEOs was better than that of the monomers mixture and each monomer. It revealed that the SEOs had synergistic effects against bacteria [22]. Synergism is the interaction between two or more antimicrobial substances, which enhanced the antibacterial effect with multiple substances. So SEOs had higher antibacterial effect than one monomer [23].

Essential oils from other plants also had antibacterial activities, such as essential oils of *Mentha citrata* Ehrh., *Artemisia annua* L. and *Citrus medica* L.. The essential oils of *Mentha citrata* Ehrh. had the significant antibacterial activity against *S. aureus* at 0.5 mg/mL [16]. The MIC of the essential oils of *Artemisia annua* L. against *S. aureus* was10 mg/mL [24]. The essential oils from fruits of *Citrus medica* L. had antibacterial activity against *S. aureus* (MIC= 1.56 mg/mL) [25]. In this study, SEOs had antibacterial activity against *S. aureu*s (MIC= 0.2 mg/mL). The lower MIC of samples means the sample had better antibacterial effect. The results of antibacterial activities showed that SEOs had better antibacterial activities than essential oils from *Mentha citrata* Ehrh., *Artemisia annua* L. and fruits of *Citrus medica* L..

*E. coli*, *S. aureus* and *P. aeruginosa* are the main pathogens causing diarrhea, urinary tract infections, skin infections and respiratory diseases, which affect human health [26, 27, 28]. In recent years, these pathogens have shown resistance to various antibiotics. Finding a safe and effective fungicide is the main goal of current studies [29, 30]. Here, SEOs had significant antibacterial properties against both Gram-positive and Gram-negative bacteria. It implied that SEOs can be used as excellent natural bacteriostatic agents to inhibit the pathogenic bacteria.

#### Antifungal effects

The antifungal activities of SEOs and monomers were evaluated using *S. cerevisiae* and *Candida albicans*. The MIC and MBC values of SEOs and monomers were summarized in Table 3 and S2 Table. The MIC results showed that it had the highest antifungal activities against *S. cerevisiae*, when the concentration of SEOs was 3.2 mg/mL. The antifungal activities of the major monomers of SEOs was tested separately, including α-pinene, α-terpineol, terpinolene and verbenone. The major monomers of SEOs showed better antifungal activities than SEOs. The results showed that α-pinene had the highest antifungal activities of SEOs against *S. cerevisiae* (MIC=0.8 mg/mL, and MBC=1.6 mg/mL). The content of α-pinene in SEOs was 31.17%. Some studies also found that α-pinene had great antifungal effect [31]. The other three main monomers, including α-terpineol, terpinolene and verbenone had high antifungal activity, which implied that these monomers of SEOs, such as α-pinene, α-terpineol, terpinolene and verbenone played major roles in antifungal activities against *S. cerevisiae* and *Candida albicans*.

**Table 3.**
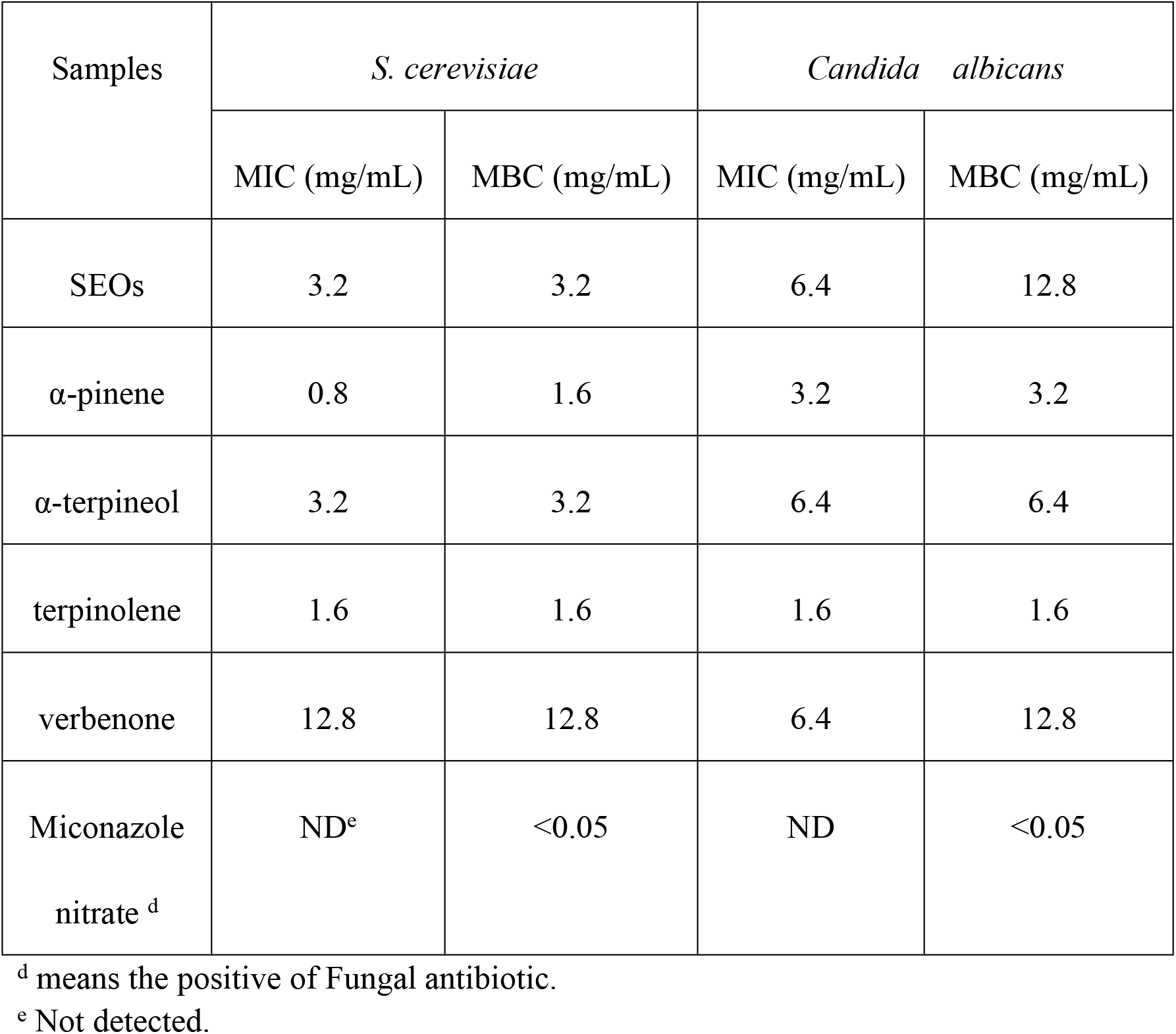
Antifungal activity (MIC and MBC) of SEOs and their major monomers.

Previous research proved that essential oils from *Euodiae Fructus* also had antifungal effects against *Candida albicans* (MIC= 25.6 mg/mL). Tthe essential oils from *Rosa* had antifungal activities against *Candida albicans* (MIC=25 mg/mL) [32, 33]. Here, SEOs had antifungal activities (MIC= 3.2 mg/mL). The results showed that SEOs had better antifungal activity than the essential oils of *Euodiae Fructus* and rose.

*Candida albicans* is the most common fungus found in gastrointestinal and genitourinary tract. The mucosa produced by *Candida albicans* causes infections in the gastrointestinal, oral and reproductive tracts, when the immunity is weak. There is no currently effective medicine for rapid clinical cure [34]. SEOs and its monomers had great antifungal activity against *Candida albicans*, which implied that SEOs could be used as natural fungicides for treating the diseases caused by *Candida albicans*.

### Antioxidant activity

#### ABTS free radical scavenging activity

ABTS scavenging rate increased with the concentration of SEOs increasing in the early stage (Fig1). ABTS scavenging rate were 37%, 54%, 75%, 84%, 90%, and 96%, when the concentrations of SEOs were 0.1 mg/mL, 0.2 mg/mL, 0.4 mg/mL, 0.6 mg/mL, 0.8 mg/mL, and 1.0 mg/mL. The results of ABTS free radical scavenging showed that the free radical scavenging rate reached 96% when the concentration of SEOs was 1.0 mg/mL. So SEOs had the high antioxidant activitiy to scavenge free radicals.

**Fig 1.**
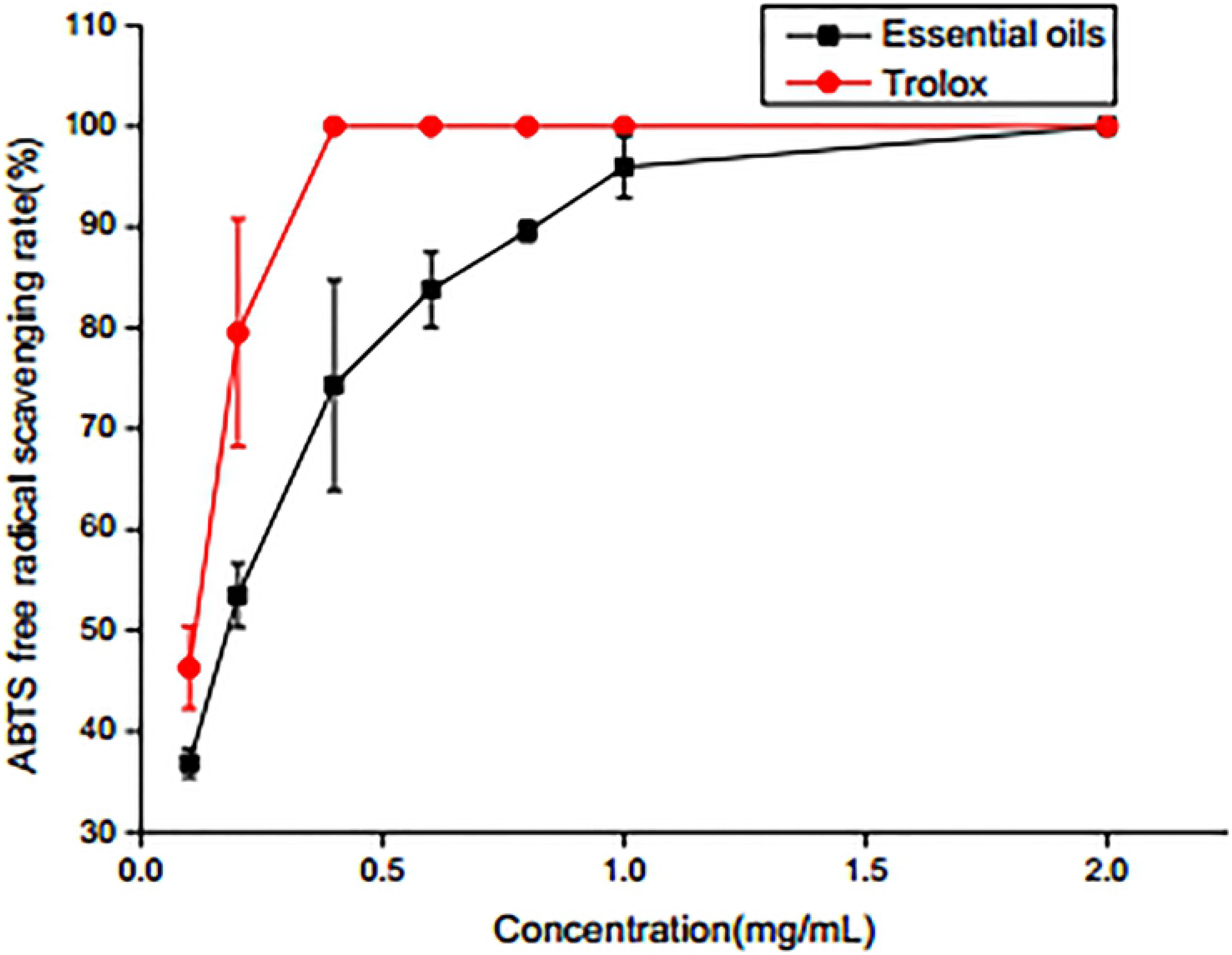
ABTS radical scavenging activity. Values are mean ± SD of three experiments. At different concentrations, it has significant antioxidant capacity (P<0.05).

#### DPPH radical scavenging activity

DPPH is a stable free radical, which has been widely used as a tool for estimating free radical scavenging activities of antioxidant. The free radical scavenging rate depended on the concentrations of SEOs in Fig2. DPPH free radical scavenging rate were 15.90%, 22.40%, 39.01%, 65.94%, 75.25%, 92.57% and 100%, when the concentrations of SEOs were 1 mg/mL, 2 mg/mL, 4 mg/mL, 6 mg/mL, 8 mg/mL, 9 mg/mL and 10 mg/mL. DPPH free radical scavenging ability of SEOs reached 100%, when the concentration of SEOs was 10 mg/mL, which is the same as that of Trolox. The results of DPPH free radical scavenging activities indicated that SEOs had high antioxidant activity. The antioxidant properties of terpenes and their derivatives were similar to those of phenolic compounds, which can scavenge free radicals by supplying hydrogen to hydrogen atoms [35]. It has been reported that α-pinene and α-celene react rapidly with peroxy radicals, resulting in a rapid termination of the oxidative chain reaction, thereby reducing the number of reactive free radicals [36].

**Fig 2.**
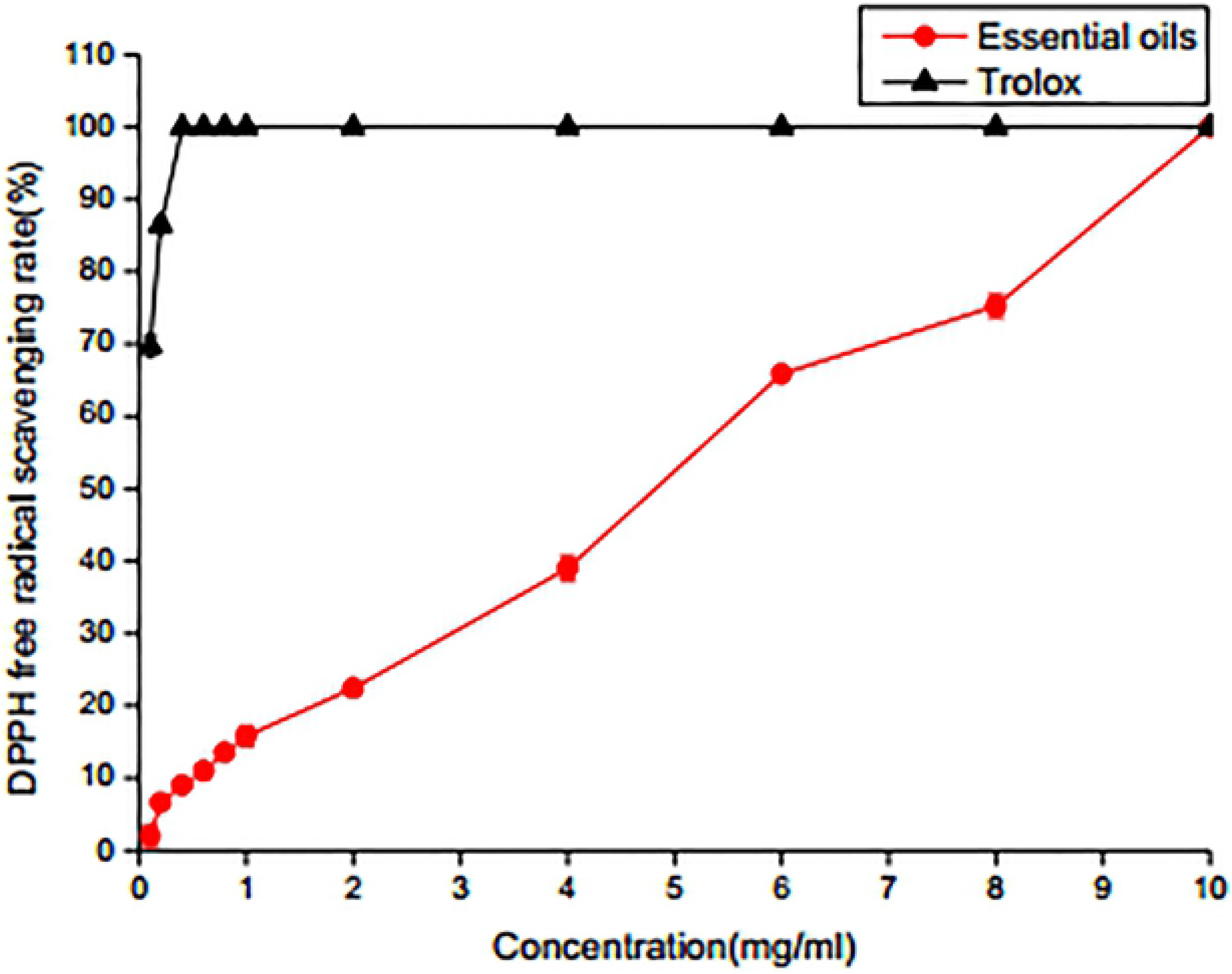
DPPH radical scavenging activity. Values are mean ± SD of three experiments. At different concentrations, it has significant antioxidant capacity (P<0.05).

#### Iron reduction ability analysis

The SEOs concentration showed an approximately linear relationship from 0.1 mg/mL to 8 mg/mL (Fig 3). The reducing power is related to the absorbance, and the reducing power increases as the absorbance increases. The results showed that the reducing ability of SEOs was related to the concentration. The higher concentration had the better reducing ability. When the SEOs reached 8 mg/mL, the reduction ability reached the highest.

**Fig 3.**
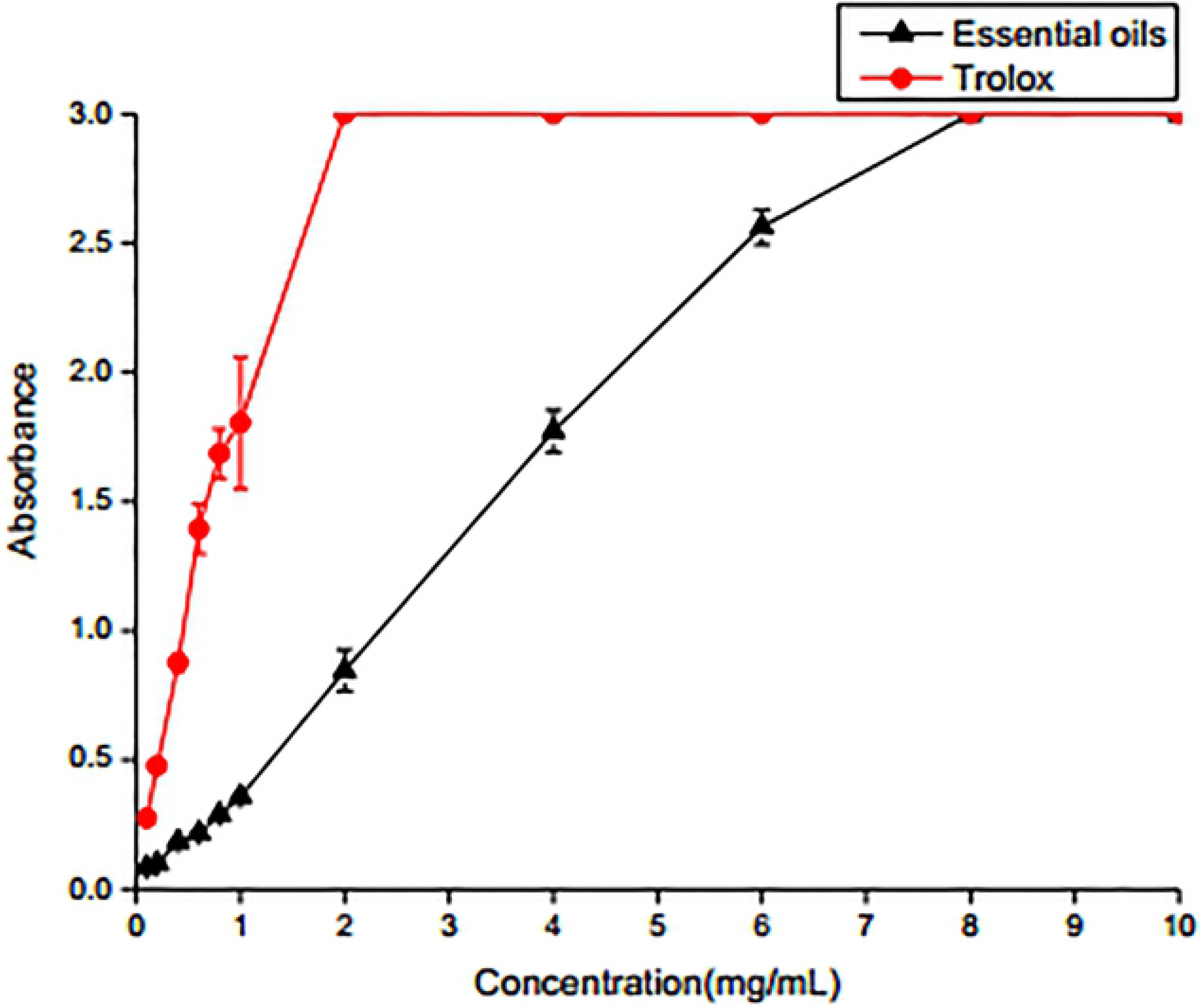
Iron ion reducing ability. Values are mean ± SD of three experiments. At different concentrations, it has significant antioxidant capacity (P<0.05).

## Conclusions

The extraction rate of SEOs was 0.4% by combining hydro-distillation and petroleum ether method. It is the first study on chemical component and biological activities of SEOs. SEOs contains 81 chemical compounds by GC-MS analysis. The main components are terpenoids, including α-pinene and α-terpineol. Through in vitro antibacterial experiments, SEOs have a high antibacterial effect on bacteria, and monomer components also have antibacterial functions. The antioxidative ability of ABTS free radicals, DPPH free radical scavenging and iron ion reduction ability was determined in vitro, which proved that SEOs had high antioxidant effect. Therefore, SEOs could be used as natural antibacterial and antioxidants agents in food, cosmetic and pharmaceutical industries.

## Acknowledgments

This study was funded by National Natural Science Funds of China (Grant No.31870201), and research funding from the Key Laboratory for Evolution of Past Life and Environment in Northeast Asia (Ministry of Education, China). We thank Jilin Teyi Food Technology Co., Ltd providing research funding and plant materials for this research.

## Author Contributions

Conceived and designed the experiments: Xinsheng Liu, Wannan Li, Bo Gao, Lu Han. Performed the experiments: Xinsheng Liu, Zian Qiao. Analyzed the data: Xinsheng Liu, Bo Gao, Lu Han. Wrote the paper: Xinsheng Liu, Xinlu Li, Lu Han.

## Supporting information

**S1Table.Antibacterial Activity (MIC and MBC) of SEOs and the major monomers.**

^a^ Mean value of absorbance.

^b^ Not detected.

^c^ mixture consists of α-pnene, α-terpineol, terpinolene, and verbenone (m: m: m: m=1:1:1:1).

^d^ Tetracycline Hcl, Bacterial antibiotic.

**S2Table.Antifungal activity (MIC and MBC) of SEOs and their major monomers.**

^e^ Miconazole nitrate, Fungal antibiotic.

